# Patterns of Saliency and Semantic Features Distinguish Gaze of Expert and Novice Viewers of Surveillance Footage

**DOI:** 10.1101/2022.01.09.475588

**Authors:** Yujia Peng, Joseph M. Burling, Greta K. Todorova, Catherine Neary, Frank E. Pollick, Hongjing Lu

## Abstract

When viewing the actions of others, we not only see patterns of body movements, but we also “see” the intentions and social relations of people, enabling us to understand the surrounding social environment. Previous research has shown that experienced forensic examiners—Closed Circuit Television (CCTV) operators—convey superior performance in identifying and predicting hostile intentions from surveillance footages than novices. However, it remains largely unknown what visual content CCTV operators actively attend to when viewing surveillance footage, and whether CCTV operators develop different strategies for active information seeking from what novices do. In this study, we conducted computational analysis for the gaze-centered stimuli captured by experienced CCTV operators and novices’ eye movements when they viewed the same surveillance footage. These analyses examined how low-level visual features and object-level semantic features contribute to attentive gaze patterns associated with the two groups of participants. Low-level image features were extracted by a visual saliency model, whereas object-level semantic features were extracted by a deep convolutional neural network (DCNN), AlexNet, from gaze-centered regions. We found that visual regions attended by CCTV operators versus by novices can be reliably classified by patterns of saliency features and DCNN features. Additionally, CCTV operators showed greater inter-subject correlation in attending to saliency features and DCNN features than did novices. These results suggest that the looking behavior of CCTV operators differs from novices by actively attending to different patterns of saliency and semantic features in both low-level and high-level visual processing. Expertise in selectively attending to informative features at different levels of visual hierarchy may play an important role in facilitating the efficient detection of social relationships between agents and the prediction of harmful intentions.

**Author Summary:** Imagine seeing a person walking toward another person menacingly on the street, we may instantly feel that some physical confrontation will happen in the next second. However, it remains unclear how we efficiently infer social intentions and outcomes from the observed dynamic visual input. To answer this question, CCTV experts, who have years of experience on observing social scenes and making online predictions of the action outcomes, provide a unique perspective. Here, we collected experts’ and novices’ eye movements when observing different action sequences and compared the attended visual information between groups. A saliency model was used to compare low-level visual features such as luminance and color, and a deep convolutional neural network was used to extract object-level semantic visual features. Our findings showed that experts obtained different patterns of low-level and semantic-level features in visual processing compared to novices. Thus, the expertise in selectively attending to informative features at different levels of visual hierarchy may play an important role in facilitating the efficient detection of social relationships between agents and the prediction of harmful intentions.

## 1. Introduction

Viewing the actions of others can reveal the intentions and social relations among people, enabling us to understand the social landscape around us. People are adept at perceiving goal-directed actions and inferring intentions from human actions. In their pioneering work, Heider and Simmel (1994) presented video clips showing three simple geometrical shapes moving around and asked human observers to describe what they saw. Almost all observers described the object movements in an anthropomorphic way, reporting a reliable impression of animacy and meaningful social interaction among the geometric shapes displayed in the decontextualized animation. However, such laboratory stimuli are limited to capture the wide range of complexity in human activities. In the everyday social environment, human behavior often involves interactions with multiple people and/or objects and is guided by sophisticated inferences about intentionality and social motivation. Although laboratory research using controlled stimuli (e.g., Heider-Simmel-type animations) has shed light on the visual processing involved in analyzing goal-oriented activities, most work has focused on how low-level visual cues, such as orientation and speed, affect social perception (e.g., Gao, McCarthy, & Scholl, 2010; Gao, Newman, & Scholl, 2009; McAleer & Pollick, 2008; Shu, Peng, Fan, Lu, & Zhu, 2018). It remains unclear how visual features at the perceptual level and semantic features at the conceptual level jointly influence how well people infer intentions in complex, real-world interactions.

We address these questions using real-life interactions recorded in videos of Closed Circuit Television (CCTV). The use of CCTV to monitor human activity in natural urban environments has become ubiquitous in societies worldwide. These systems typically employ a set of cameras deployed around complex urban geography. The videos recorded by the cameras are routinely monitored by CCTV human operators in real-time to identify the presence of hostile intentions so as to allow a preemptive response that minimizes the consequences of these intentions (Wallace, & Diffley, 1998), as well as to obtain evidence when events do occur.

Surveillance CCTV videos usually contain a large amount of visual information coupled with the high complexity of human activities. Hence, CCTV operators, who have acquired extensive experience in the visual analysis of human actions in real-world scenes, likely adopt efficient strategies in information processing. Previous studies have compared CCTV operators to novices when performing the task of judging harmful intent from CCTV videos. Most studies have found group differences between CCTV operators and novices in brain activity and in the ability to recognize and predict harmful intention (Gillard et al., 2019; Grant & Williams, 2011; Howard et al., 2009; Petrini, McAleer, Neary, Gillard, & Pollick, 2014; but also see Troscianko, Holmes, Stillman, Mirmehdi, Wright, & Wilson, 2004). Hence, studying the differences in information processing between experienced CCTV operators and novices provides a unique window to unveil efficient strategies acquired by human experts through extensive learning.

What learning-induced processes lead to the differences in neural mechanisms and enhanced behavioural performance of CCTV operators for video analysis and intention identification? In the perceptual learning literature, there is ample evidence to support learning with intensive visual experiences modulating the connection weights between basic visual channels and decisions (e.g., Dosher & Lu, 1998), which enable efficient feature selection for a trained task. For CCTV operators with years of visual experience in action perception, we hypothesize that this extensive experience in viewing surveillance footage likely promotes selective processes for visual information by actively attending to different contents in visual inputs.

Humans do not examine the visual environment in a passive manner as does a camera taking pictures; rather, we actively sample the visual input through brief fixations interspersed with gaze shifts. During a period of stable fixation, the information at the central gaze is analyzed in fine detail using foveal vision. At the same time, peripheral analysis is carried out to select the next fixation location for a gaze shift. Recent studies have shown significant differences between central and peripheral vision in the analysis of human body movements (Thurman & Lu, 2013, 2014), showing that configural cues based on the spatial arrangement of joint trajectories dominate visual processing in central vision, whereas local motion and orientation cues interact with spatial cues to influence action perception in the periphery. In addition, studies investigating surveillance videos have provided evidence that experienced CCTV operators, relative to novices, produce different goal-directed eye movement patterns when viewing surveillance video, and show greater consistency in eye-movement tracking patterns (Howard, Troscianko, Gilchrist, Behera, & Hogg, 2013; Roffo, Cristani, Pollick, Segalin, & Vittorrio, 2013). Although these studies have analyzed eye-movement characteristics associated with expertise and behavior, it remains unknown what stimulus content in surveillance videos drives the active selection of gaze shifts when viewing people’s activities with the goal of identifying intentions and whether CCTV operators develop different information-seeking strategies than novices.

To understand the difference in visual content attended by CCTV operators versus novices, we conducted two computational analyses. These analyses focused on investigating how both low-level visual features and object-level semantic features contribute to attentive gaze patterns associated with the two groups of participants. The first analysis adopted a saliency model (Itti, Koch, & Niebur, 1998) to characterize low-level visual features attended by gaze.

The saliency model decomposes visual inputs into a set of topographic feature maps, such as motion, luminance, color, texture, and orientation (Treisman & Gelade, 1980). All feature maps feed, in a purely bottom-up manner, into a master “saliency map,” which topographically codes for local conspicuity over the entire visual scene. Different spatial locations then compete for saliency within each map, such that only locations which locally stand out from their surround can persist. Specifically, the saliency model can compute scores reflecting the degree of gaze-centered regions capturing bottom-up visual attention in the video frames.

The second computational analysis applies a deep convolutional neural network (DCNN), AlexNet (Krizhevsky, Sutskever, & Hinton, 2012), to extract object-level semantic features from gaze-centered regions of visual inputs (Kriegeskorte, 2015). In recent years, DCNNs have provided groundbreaking results in a range of visual tasks in which the networks have shown comparable performance to human observers (Krizhevsky, Sutskever, & Hinton, 2012). DCNNs usually build on a multi-layer architecture. For example, AlexNet contains eight layers (Güçlü & van Gerven, 2015; Zeiler & Fergus 2013). This network architecture is consistent with the hierarchical structure of the visual system in human brains. The architecture enables DCNNs to cope with nonlinearity and complex visual tasks. In addition to having a similar architecture, the inner representations of DCNNs have also been found to capture neural similarities in brain activities for different visual inputs (Kriegeskorte, Mur, & Ruff, 2008; Cichy, Khosla, Pantazis, Torralba, & Oliva, 2016). For example, the later layers of DCNNs have been found to reflect neural activities in high-level visual cortex for object recognition (Yamins, Hong, Cadieu, & DiCarlo, 2013; Yamins et al., 2014). Hence, activities in later layers (e.g., fully-connected layer) of DCNNs appear to capture abstract features crucial to visual knowledge and scene semantics.

Together, the Saliency model and the DCNN model provide complementary analyses for assessing how CCTV operators and novices use various features extracted from different levels of visual hierarchy. If low-level visual saliency cues have a greater impact on capturing attention and driving the inference of intentions, we would expect to find a difference in visual saliency from the gaze-centered stimulus regions between CCTV operators and novices. On the other hand, if CCTV operators differ from novices primarily in the use of semantic features extracted by high-level visual processing, the DCNN may be able to capture the group differences. Or CCTV operators show differences from novices in the analysis of features at both processing levels. In addition, we examine the inter-subject correlation of visual features attended by CCTV operators and novices. If the expertise of CCTV operators leads to shared strategies that emerged from rich experience in analyzing surveillance footage, we would expect to find greater inter-subject correlation of visual features attended among CCTV operators than for novices. Finally, we examine how different action categories interact with information-processing differences between CCTV operators and novices. We hypothesize CCTV operators may be able to extract visual information signalling antisocial intentions more efficiently (e.g., actions ending with fights or confrontation) while showing less or even no differences from novices for actions involving benign intentions.

## 2. Methods

### 2.1 Participants

Eleven CCTV operators (3 female, aged 21-53 years, *M* = 36.3, *SD* = 10.1) and ten novices (2 female, 8 male, aged 28-43 years, *M* = 33.8, *SD* = 6.0) were recruited to participate in the eye-tracking experiment. The ‘operator’ participants were all employed to monitor CCTV when the experiment was conducted and had an average of 4.5 years of working experience as a CCTV operator (*SD* = 3.0, range 0.4-12 years), and viewed CCTV an average of 10 hours (range 8-12 hours) per day. The ‘novice’ participants were defined as individuals with no CCTV surveillance or security experience. The age of the operators and novices were matched (independent t-test, *t*(19) = 0.875, *p* = 0.392). CCTV operators were recruited from CCTV control rooms and user groups. Novices were recruited from the community through advertisements.

Each participant read and signed a Consent Form that described their participation in the experiment and the use of the data collected. All participants were free to leave the study at any time. Ethical approval for all phases of the study was obtained from the UK Ministry of Defense Research Ethics Committee. Participants were paid for their time and travel expenses to attend the experiment at BAE Systems, Advanced Technology Centre, Bristol.

### 2.2 Stimuli and procedure

Videos of street scenes with human actions recorded by CCTV were selected from originally over 800 h of CCTV footage obtained of urban scenes in the UK. Four paid research assistants with no prior CCTV experience screened the corpus of video material and identified CCTV clips that resulted in physical aggression (and therefore included hostile intent), which were labeled as the ‘Fight’ clips. Control scenes were chosen for the ‘Confrontation’, ‘Playful’, and ‘Neutral’ categories and were matched to the Fight clips in several respects: location, time of day, and the number of people in each display. A total of 36 videos were generated for the four action categories, with nine videos in each category (for more details, see Petrini, McAleer, Neary, Gillard, & Pollick, 2014). Each video lasted 16s with a frame rate of 25 fps, yielding a total of 400 frames, with the image size of 576 × 480 pixels in a visual angle of 22.5°×19°. The fight videos showed 16 sec of aggressive behavior prior to a violent incident. Eye movements were recorded when participants viewed these 36 videos. Eye movement data were collected using an ASL Eye-Trace6 system with a sample rate of 50Hz and accuracy approximately 1 degree across the visual field. For each participant, we extracted square image patches from each video frame with a window size of 75 × 75 pixels centered on a gaze-fixation location in each frame. The size gaze-contingent window (approximately 3°× 3°) is larger than the standard estimate of the foveola, which is roughly 0.5-1.0° in diameter (Boff & Lincoln, 1988). We also examined different window sizes of 38 × 38 (1.5°× 1.5°) and 150 × 150 (6°× 6°) and got similar results. Certain video frames had missing eye-tracking data, with an average of 1.1% (i.e., 4.6 frames among 400 frames).

### 2.3 Computational analyses

Two computational models were used to extract visual features from the raw videos of surveillance footage: a saliency model and a deep convolutional neural network model. An illustration of the procedure is shown in Figure 1. The saliency model was adopted to capture low-level image features that attract attentive gaze, and the DCNN model was adopted to capture object-level features that capture semantic information in attended visual scenes. For frames with missing eye tracking data, blank image patches were extracted and zeros were used for computational models.

**Figure 1:**
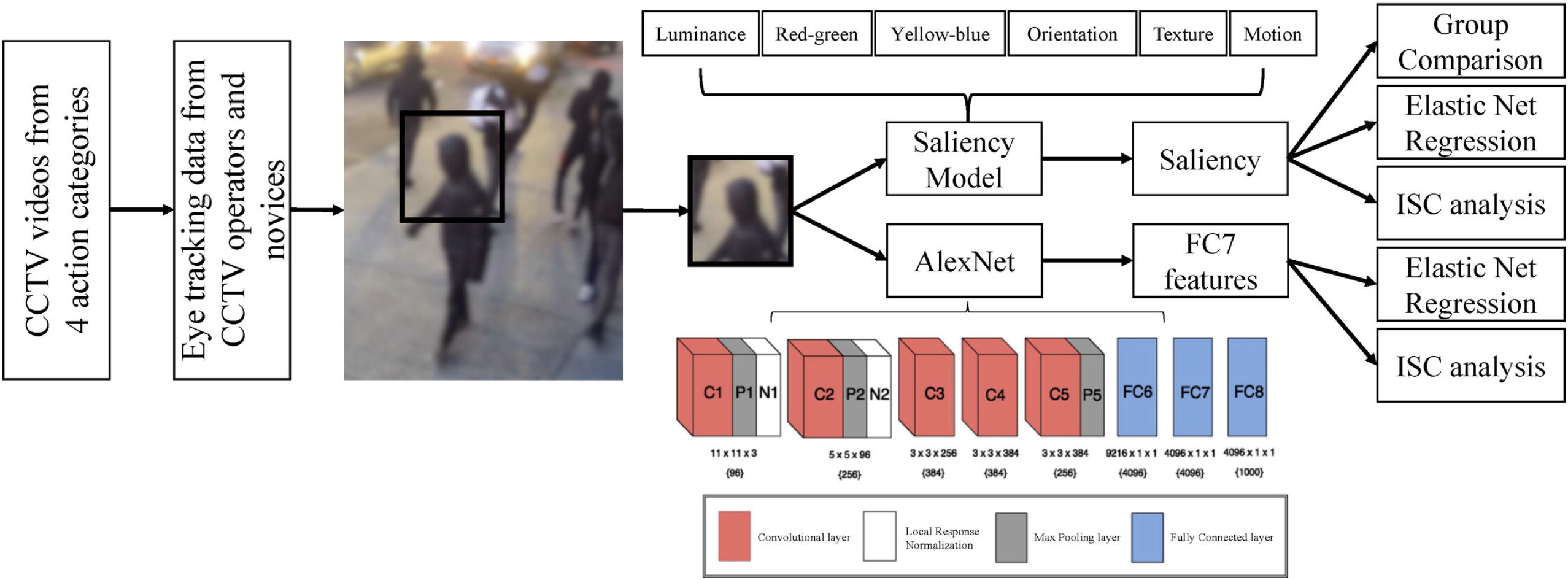
Procedures for feature extraction and comparison. A square image patch centered on coordinates of fixations was extracted from each image frame and these were fed into models as input. The saliency model extracted saliency features, which were used to derive a saliency index to compare across groups. The extracted saliency features were also entered into an elastic net regression model for decoding CCTV operators from novices. The AlexNet extracted fully-connected layer features as inputs for entry to an elastic net regression model for the decoding purpose. Inter-subject correlation (ISC) indices based on saliency features and DCNN features were compared between groups. Note, the blurred image frame was selected for demonstration and was not from the real experimental materials.

#### 2.3.1 Saliency model and saliency index

We adopted the saliency model by Itti and colleagues (Itti, Koch, & Niebur, 1998) to analyze the influences of low-level visual cues on gaze patterns. The saliency model processes visual input in a bottom-up manner and does not capture high-level visual features associated with objects or people. As shown in Figure 2, image features were extracted from each image frame through six processing channels: luminance, color (red-green and yellow-blue), orientation, texture, and motion. Luminance and color maps were calculated based on the Derrington-Krauskopf-Lennie (DKL) color space (Derrington, Krauskopf, & Lennie, 1984) using long, medium, and short cone response filters. Luminance maps were computed as the sum of long and medium cone responses. Colors were defined as the difference between long and medium for the red-green colormap and (Long + Medium) - Short for the yellow-blue colormap. The orientation map was created by applying a series of Gabor filters to the grayscale image to detects line-segment edges. The texture map was created by applying a series of Laplacian of Gaussian (LoG, or Mexican hat) filters of different spatial sizes proportional to the grayscale image size. The optical flow map was estimated using an orientation tensor (Farnebäck, 2000), which processes the current and the previous image frame to detect shifts in location of each pixel in temporally neighboring frames. Only vector magnitude was used to represent optical flow magnitudes (i.e., motion speed) without the consideration of motion directions.

**Figure 2:**
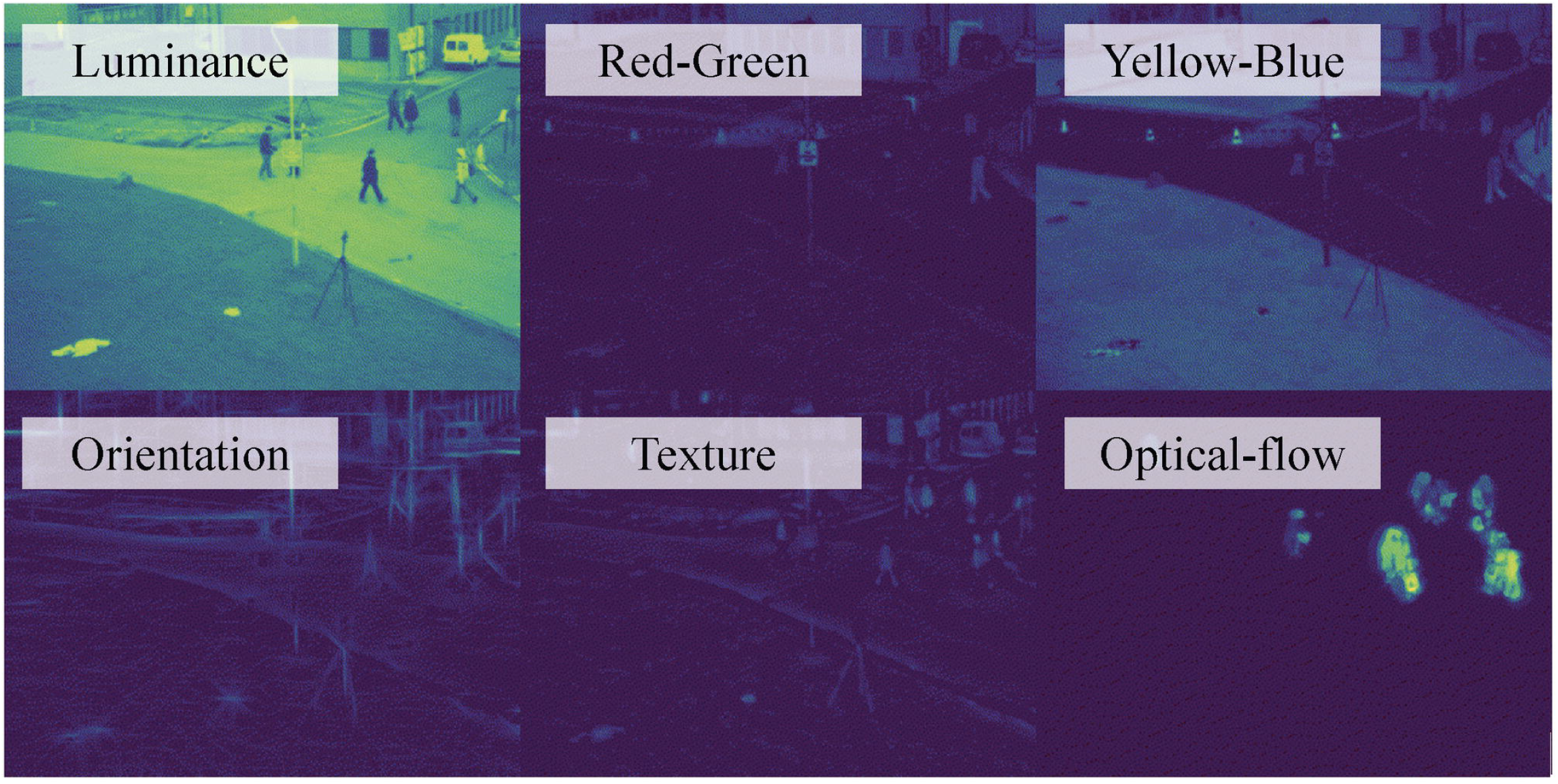
Image features were extracted from each image frame through six processing channels in the saliency model. Note, the image frame was selected for demonstration and was not from the real experimental materials.

For each image frame, a pyramid stack was created for each feature channel that included the original size of the source image patch (75 × 75 pixels), half size, quarter size, and eighth size scales of the image. Gaussian blur was applied before each downsampling operation.

Pyramids achieved the result of increasing the receptive field during the activation step. A center-surround activation step was implemented as a combination of Laplacian-Gaussian convolution and Gaussian blur, and was applied to each map iteratively for a total of five passes of the convolution kernels. All convolution kernels used to extract feature maps during activation were applied to the source image. A temporal buffer of the pyramid stacks was created to keep track of the previous two frames and the current frame. The buffer was used to record previous feature channel weights and final saliency maps, acting as a weighted memory for past salient regions. For each new frame, past saliency images were weighted by half of their current values.

After extracting and processing the pyramid stacks, feature maps have different value ranges. The normalization step obtains a common scale across all feature maps so that they can later be combined into a single saliency map. For normalization, each feature map within a pyramid stack was scanned to enhance the contrast between salient and non-salient regions. The sum of feature values for the salient regions was used to compute normalization factors to be applied to the set of feature maps. The normalization factors for the current source image were weighted by the normalization factors stored in the temporal buffer, and the current maps were then scaled according to the weighted normalization factors. The weighted normalization factors used for subsequent analysis were exported for each image frame to assess the relative contribution of each feature channel.

To calculate the final saliency map for an image frame, first, the feature maps were compressed into a single map by summing across pyramid levels for each channel and dividing by the size of a pyramid stack, yielding an intermediate saliency map for each feature channel. Secondly, the intermediate saliency maps per feature channel were summed to create a single saliency map for the current image frame. The saliency map for the current image frame was then combined with the weight-decayed saliency maps from previous two frames and the current frame in the temporal buffer to obtain a final saliency map. Lastly, the final map was processed with a logistic activation function that increases the contrast between salient and non-salient regions, which was also exported for analysis. Using the final saliency maps from a sequence of image frames, a *saliency index* was calculated by computing the average saliency values within a gaze-centered region in the saliency maps.

#### 2.3.2 Saliency analysis

To examine whether CCTV operators and novices’ gaze patterns are impacted differently by low-level saliency cues, we conducted repeated-measures ANOVAs for each action category to examine the group difference between CCTV operators and novices on the saliency index and the six feature dimensions. We hypothesize that if the visual contents attended by CCTV operators in their gaze fixations differ from the visual information captured by novices in terms of salient low-level features (e.g., luminance or motion features), we would expect to observe a group difference between the two groups of participants in the saliency index obtained from gaze-centered regions derived from their eye movement patterns.

Additionally, we concatenated six saliency features to form a multidimensional saliency vector to train a machine learning classifier based on elastic net regularization (Tibshirani, 1996; Zou & Hastie, 2005). The classifier was trained to differentiate CCTV operators and novices based on the attended low-level saliency information. Specifically, the concatenated feature vectors were entered as predictors to the generalized linear model (GLM) to classify CCTV operators and novices. The classifier used the elastic net regularization to favor the selection of a small number of important features that help predict the class labels. The elastic net regularization has a free parameter, α, controlling the weight between a lasso (L1) and a ridge (L2) regularization. We set an alpha value of 1 to favor a smaller number of features. We also used a parameter value of 0.9 and 0.8 that yielded similar results. The model was trained in a leave-one-out manner with 21 iterations. Specifically in each iteration, 20 participants were randomly selected for training, and the remaining one participant was used for testing to let the classifier determine whether this testing participant was a CCTV operator or a novice. Classification accuracy was averaged across all 21 iterations.

To reflect the online processing with cumulative information over time, for each video, features were concatenated across a set of non-overlapping cumulative temporal windows with a step-size of 50 frames (i.e., concatenating features of frames 1 to 50, frames 51 to 100, …, and frames 351 to 400), yielding eight chunks of feature vectors. Using cumulative frames by concatenation takes into consideration the temporal dependency in action videos. We explored a set of temporal cumulation windows because critical events occurred at different time points for different surveillance footage.

Since the most informative signal that differentiates CCTV operators and novices may emerge at different time points, the *maximum* classification accuracy over temporal cumulation windows was used for each video as the decoding accuracy of the classifier. For example, for confrontation video No.1, the maximum classification accuracy may arise from early frames of the video with cumulating frames 1-50, while confrontation video No. 2 may reveal the maximum classification accuracy from a different temporal window of frames 51-100. If operators and novices attended to systematically different low-level features captured through the saliency model, we would expect that the classifier should show above chance-level accuracy in differentiating operators and novices. Furthermore, if attentive features were influenced by the nature of intention underlying the observed actions, the classifier accuracy may vary depending on the presence or absence of harmful intentions. Bonferroni multiple-comparison correction was applied to the statistical testing of decoding accuracy.

Furthermore, to examine whether operators or novices consistently attend to information with high saliency, inter-subject correlations were calculated for the operator group and the novice group separately. For image patches centered at gaze fixations in each frame, a saliency vector was extracted from all six feature maps. To transform features onto a common scale and remove outliers, z-score normalization was applied for each feature channel across all videos and subjects. The similarity of gaze-centered saliency between a pair of participants was computed as the correlation of concatenated saliency vectors over time for each video (i.e., each video yields a 2400-element-long vector coming from 6 features of 400 frames). Higher similarity values indicate that two participants attended to regions with a similar degree of visual saliency. For each video, the inter-subject correlation (ISC) was defined as the average similarity value across all pairs of participants within the operator group and within the novice group.

#### 2.3.3 Deep convolutional neural network model and analysis

In the DCNN analysis, due to the high similarity of objects involved in consecutive frames, one frame out of every ten frames was sampled as inputs into models. Thus, the original 400 frames of surveillance footage were downsampled to 40 frames to reduce computational demands. To investigate the group difference on a semantic level, we adopted a pre-trained DCNN, AlexNet, to extract object-level features. AlexNet contains five convolutional layers and three fully connected layers. For each image patch centered at the gaze fixation, we extracted the activations from the penultimate layer, fully connected layer 7 (FC7, containing 1*4096 units), a layer just before the decision layer for object categorization in AlexNet. For each video (36 videos in total), features of image patches centered on fixations were extracted from the penultimate layer (i.e., FC7) of AlexNet. Each gaze-centered image patch yields a feature vector in a size of 1 by 4096. Similar to the analysis approach used for the saliency model, to reflect the online processing with cumulative information over time, for each video, features were concatenated across a set of cumulative frame windows with a step-size of 5 frames (i.e., concatenating features of frames 1 to 5, frames 6 to 10, …, and frames 36 to 40). Because the DCNN features were downsampled by a factor of 10, current windows with 5 frames match what was used for the saliency model. The classifier with elastic net regularization was applied on the DCNN features to differentiate visual information attended by CCTV operators and novices.

Training and testing procedures were the same as the classifier with saliency features. If operators and novices attended to systematically different object-level features captured through the DCNN model, we would expect the classifier to show the above chance-level accuracy in differentiating operators and novices.

Similar to the saliency analysis, to examine whether operators or novices consistently attend to object-level information extracted by DCNN, inter-subject correlations were calculated for the operator group and the novice group separately. First, FC7 features of all gaze-centered image regions were concatenated across time by video. ISC was calculated as the correlation coefficients of concatenated feature vectors between pairs of operators, or between pairs of novice participants. This procedure was repeated for each of the 36 videos, respectively.

## 3. Results

### 3.1 Saliency analyses

#### 3.1.1 Saliency indices of CCTV operators and novices

Figure 3 depicts the saliency index for image patches in the gaze fixation areas as a function of video time for the two groups of participants, CCTV operators and novices. Mixed ANOVA models with participant group as a between-subjects factor and time as a within-subjects factor were conducted on the saliency index for each of the four action categories. For all four types of actions, we found significant main effects of time (Fight, *F*(7,13) = 12.30, *p* < .001, η_*p*_^2^ = .869; Confrontation, *F*(7,13) = 69.33, *p* < .001, η_*p*_^2^ = .974; Playful, *F*(7,13) = 33.18, *p* < .001, η_*p*_^2^ = .947; Neutral, *F*(7,13) = 8.50, *p* = .001, η_*p*_^2^ = .821). This result suggests that visual saliency changes dynamically across video frames, which is consistent with the complex nature of visual contents in the surveillance footages. However, no action type showed a main effect of participant group, revealing that image patches attended by CCTV operators and novices do not show significant differences in terms of visual saliency (Fight, *F*(1,19) = 2.07, *p* = .166, η_*p*_^2^ = .098; Confrontation, *F*(1,19) = .949, *p* = .342, η_*p*_^2^ = .048; Playful, *F*(1,19) = 2.77, *p* = .113, η_*p*_^2^ = .127; Neutral, *F*(1,19) = 0.087, *p* = .771, η_*p*_^2^ = .005). The lack of group difference in the saliency index of gaze-centered regions suggests that low-level visual saliency is not a strong cue to differentiate the information used by CCTV operators from that used by novices. The two-way interaction effect between time and participant groups was not significant for any of the action types (Fight, *F*(7,13) = 1.60, *p* = .221, η_*p*_^2^ = .462; Confrontation, *F*(7,13) = .615, *p* = .735, η_*p*_^2^ = .249; Playful, *F*(7,13) = .433, *p* = .865, η_*p*_^2^ = .189; Neutral, *F*(7,13) = .925, *p* = .518, η_*p*_^2^ = .333).

**Figure 3:**
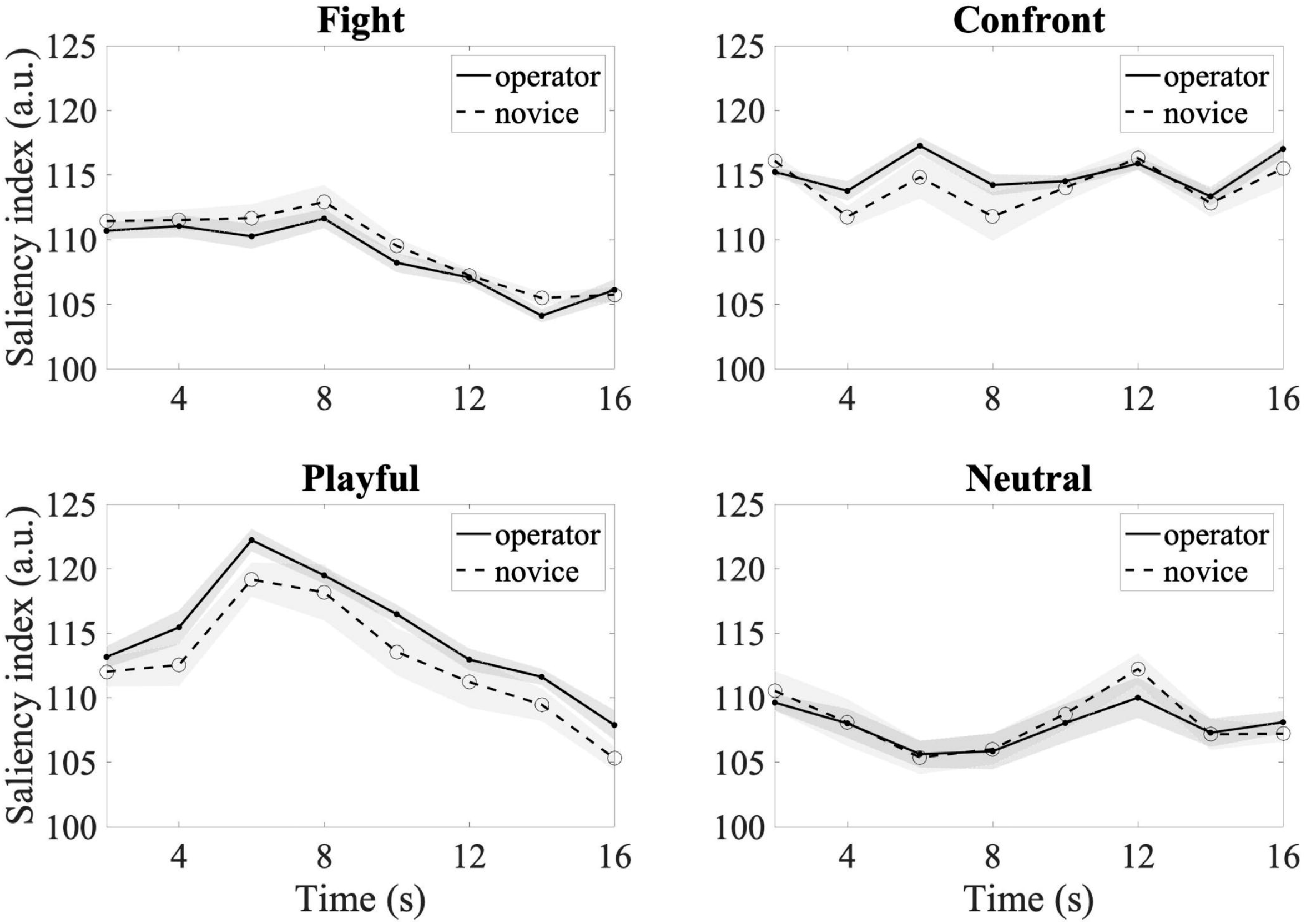
Saliency indices of the operator and novice groups over time for four types of actions. Each data point corresponds to the averaged saliency index every 2 seconds. Shaded areas indicate standard error.

To examine the robustness of the results and to make sure the effects were not driven by a difference in the proportion of missing frames between operators and novices, we reran the analysis after removing missing frames. Similar results were found for all four action categories. For all four types of actions, we found significant main effects of time (*p*^2^s < .001), but neither the main effect of participant group or the two-way interaction effect between time and participant groups were significant (*p* > 0.05).

#### 3.1.2 Saliency feature decoding of CCTV operators and novices

We next conducted a multivariate classification analysis to investigate whether CCTV operators and novices could be decoded based on patterns of saliency features. Saliency feature vectors were concatenated across frames and were used to train classifiers to recognize visual information attended by operators or by novices. As shown in Figure 4a, all four actions reached decoding accuracy that was significantly greater than the chance level 0.5. Four one-sample t-tests were carried out and each tested against a Bonferroni-adjusted alpha level of 0.00625 (0.05/8 for eight time chunks). Fight videos reached the highest averaged classification accuracy (*M* = 0.70, *SD* = 0.09) (*t*(8) = 6.92, *p* < 0.001), but was not significantly different from other action categories (Confrontation, *M = 0.67, SD = 0.05, t(8) = 9.39, p < 0.001*; Playful, *M = 0.68, SD = 0.10, t(8) = 5.28, p < 0.001*; Neutral, *M = 0.70, SD = 0.10, t(8) = 5.59, p < 0.001*). We also reran the analysis after removing videos with more than 10% of missing eye-tracking data. Similar decoding results were found for all four action categories (Fight, *M = 0.67, SD = 0.14, p = 0.007*; Confrontation, *M = 0.68, SD = 0.09, p < 0.001*; Playful, *M = 0.70, SD = 0.14, p = 0.002*; Neutral, *M = 0.63, SD = 0.08, p = 0.001*).

To examine the contribution of six saliency features in the decoding algorithm, we calculated the proportion of features being selected by the elastic net regression model. As shown in Figure 4b, optical-flow motion information was the most frequently selected feature to differentiate operators from novices (*M* = 0.25, SD= 0.005), followed by texture (*M* = 0.18, SD= 0.003), orientation (*M* = 0.16, SD= 0.004), luminance (*M* = 0.16, SD= 0.003), and yellow-blue color (*M* = 0.15, SD = 0.003).

**Figure 4:**
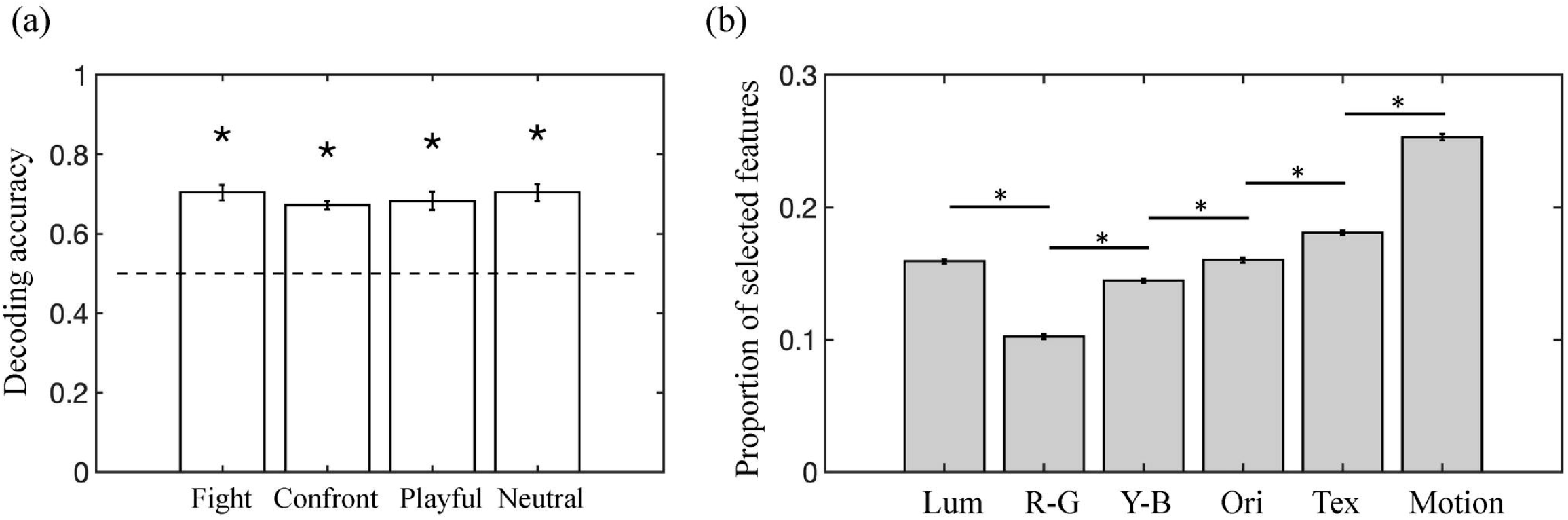
*(a) Decoding accuracy based on saliency features of discriminating CCTV operators from novices. Error bars indicate SDs* of accuracy from leave-one-out iterations. *Asterisks indicate significantly greater than the chance level tested against a Bonferroni corrected alpha level. (b) Proportions of saliency features selected in the elastic net regression decoding analysis. From left to right, the six feature dimensions represent luminance, red-green color, yellow-blue color, orientation, texture, and optical-flow motion information. Error bars indicate SEs* of CCTV operators and novices. *Asterisks indicate significant differences between feature dimensions*.

#### 3.1.3 Inter-subject correlation (ISC) of saliency index

To examine whether operators or novices consistently attend to information with high saliency, inter-subject correlations were calculated for the operator group and the novice group separately. Saliency ISC of each action category within the groups are shown in Fig 5a. A repeated-measure ANOVA was conducted with groups and action categories as within- and between-subject factors. The ANOVA showed a significant main effect of the participant group, *F*(1,32) = 9.38, *p* = 0.004, *η*_*p*_^*2*^ = .227, resulting from greater inter-subject correlation among experienced operators than among the novice group. The main effect of action categories was not significant, *F*(1,3) = 1.86, *p* = 0.157, *η*_*p*_^*2*^ = .148. The two-way interaction between groups and action categories was not significant, *F*(3,32) = 0.31, *p* = 0.822, *η*_*p*_^*2*^ = .028. To further understand the main effect of group, the ISC of each action category was then compared between operators and novices. Playful actions showed a significant simple main effect of group difference, *F*(1,32) = 4.30, *p* = 0.046, *η*_*p*_^*2*^ = .118 None of the other three action categories reached a significant group difference on the ISC of saliency index (Fight, *F*(1,32) = 1.15, *p* = .292, *η*_*p*_^2^ = .035; Confrontation, *F*(1,32) = 3.77, *p* = .061, *η*_*p*_^2^ = .105; Neutral, *F*(1,32) = 1.08, *p* = .307, *η*_*p*_^2^ = .033). To examine the robustness of the results, we reran the analysis with missing data removed. Videos clips with more than 10% of the data points missing were removed from analysis. Similar results were found as above after removing missing data. The main effect of the participant group was significant (*p* = 0.011), showing greater saliency ISC among experts than novices. The main effect of action categories (*p* = 0.810) and two-way interaction between groups and action categories was not significant (*p* = 0.328).

Additionally, to examine how the group difference on the consistency in attending saliency features emerges over time, we examined the dynamic change of ISC for every two seconds in time, yielding eight frame time chunks. Saliency ISC of CCTV operators were compared to novices at each time point. As shown in Fig. 5b, none of the group differences survived Bonferroni correction (Bonferroni-adjusted alpha level = 0.00625), except that a marginally significant effect (*p* = 0.009) was found in the window between 10 to 12s.

**Figure 5:**
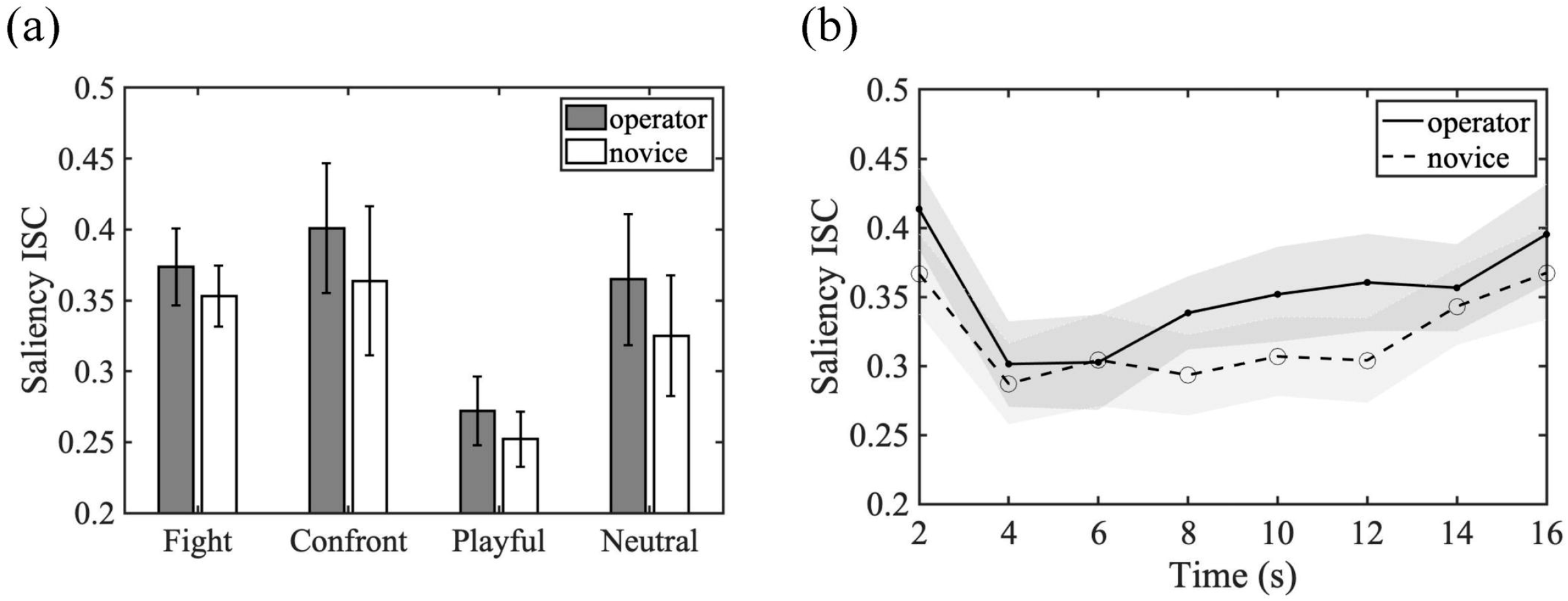
Saliency inter-subject correlation (ISC) results. (a) Saliency ISC of each action category within the CCTV operator group and within the novice group, calculated by averaging pairwise correlations of saliency features concatenated over time for each video. Error bars indicate SEs across videos in each action category. (b) Saliency ISC integrated within every 2s time windows for CCTV operators and novices. Shaded areas indicate SEs.

### 3.2 DCNN feature analysis

#### 3.2.1 DCNN decoding of CCTV operators and novices

Features extracted from DCNN were used to train a classifier to recognize visual information attended by operators or by novices. One sample t-tests were carried out for each action category and tested against a Bonferroni-adjusted alpha level of 0.00625 (0.05/8 for eight time chunks and 4 action categories included in the analysis). As shown in Fig. 6, all except neutral actions reached a classification accuracy that was significantly above the chance level (i.e. 50%) (Fight: *M* = 0.71, *SD* = 0.10, *t*(8) = 6.49, *p* < 0.001; Confrontation: *M* = 0.74, *SD* = 0.09, *t*(8) = 7.90, *p* < 0.001; Playful: *M* = 0.71, *SD* = 0.15, *t*(8) = 4.25, *p* = 0.003; Neutral: *M* = 0.67, *SD* = 0.14, *t*(8) = 3.589, *p* = 0.007). We also examined the robustness of the decoding results by removing two subjects whose gaze sequences yielded excessive missing data. The decoding results with 19 subjects showed similar results as before. However, only fighting actions reached a significant classification accuracy (*M* = 0.70, *SD* = 0.10, *p* < 0.001), surviving a Bonferroni-adjusted alpha level, where as other action categories did not survive the multiple comparison correction (Confrontation: *M* = 0.67, *SD* = 0.15, *p* = 0.009; Playful: *M* = 0.60, *SD* = 0.10, *p* = 0.024; Neutral: *M* = 0.65, *SD* = 0.14, *p* = 0.013).

**Figure 6:**
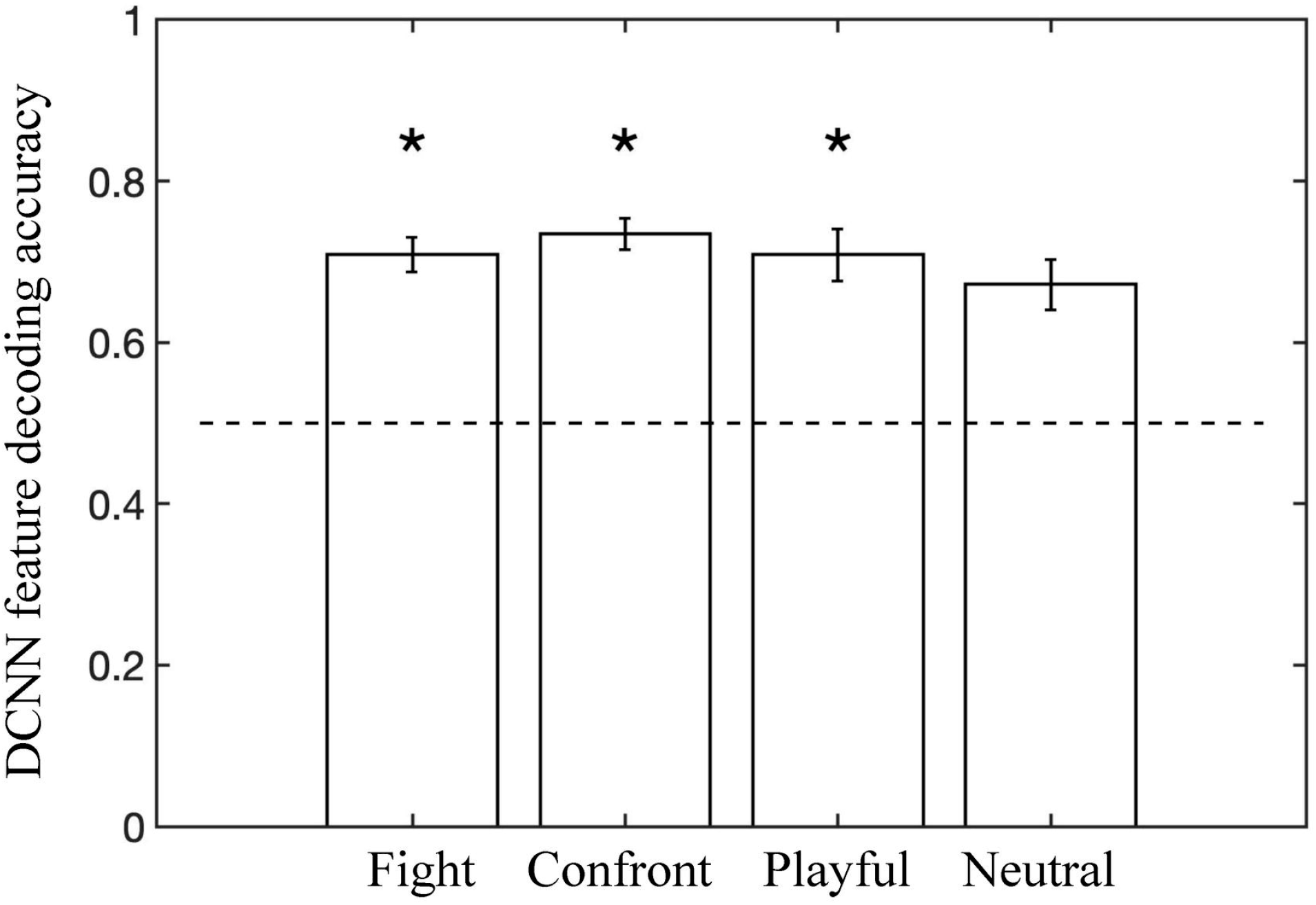
Decoding accuracy based on fully-connected layer CNN features on discriminating CCTV operators from novices. Fight, confrontation, and playful action categories yielded above-chance decoding accuracy after Bonferroni correction. Error bars indicate SEs of accuracy from leave-one-out iterations. Asterisks indicate significantly greater than chance level tested against a Bonferroni corrected alpha level.

These results suggest that the DCNN features for gaze-centered regions were able to classify CCTV operators from novices for actions governed by potentially harmful intentions, especially for fighting actions where physical violence ensued after completion of the clip. In contrast, for actions with less clear intentions in the neutral condition, the decoding based on DCNN features was weak as the accuracy did not survive multiple-comparison correction.

#### 3.2.2 Inter-subject correlation of DCNN features

To examine whether operators show more consistency in attending to similar DCNN features than novices, we conducted inter-subject correlation analysis. As shown in Fig. 7a, using DCNN features, ISCs averaged across nine videos in each action category were compared between CCTV operators and novices. A repeated-measure ANOVA showed a significant main effect of the participant group, *F*(1,32) = 26.26, *p* < 0.001,*η*_*p*_^2^ = 1.0, resulting from higher inter-subject correlation among experienced operators than among the novice group. The main effect of action categories was not significant, *F*(1,3) = 2.31, *p* = 0.095, *η*_*p*_^2^ = .53. The two-way interaction between groups and action categories was not significant, *F*(3,32) = 1.56, *p* = 0.219, *η*_*p*_^2^ = .37. To test the robustness of the inter-subject correlation of DCNN features, we removed gaze sequence data with more than 10% of missing data and got similar results as before.

To examine how the group difference on the consistency in attending DCNN features emerges over time, we further examined the dynamic change of ISC for every two seconds in time, yielding eight time chunks. As shown in Fig. 7b, CCTV operators and novices showed significant group differences both at the very beginning of videos (i.e., 0-2s after video onsets, *p* < .001, tested against a Bonferroni-adjusted alpha level = 0.00625) and during the latter half of video displays (i.e., from 8 s to 10 s, *p* = .002, and from 10 s to 12 s, *p* = .004, after video onsets). The *p* values for 6 s to 8 s (*p* = 0.018) 10 s to 12 s (*p* = 0.019) were less than 0.05 but did not survive the Bonferroni-adjusted alpha level. This indicates that CCTV operators showed more consistency in attending DCNN features than novices even at the onset of videos. This result suggests that operators may share some potential strategies to capture certain high-level semantic information about surveillance footages at the beginning of videos. Additionally, this result indicates that experienced operators showed greater consistency when attending to information with semantic features captured by DCNN model for an important time period later in the videos, which is critical for the recognition and prediction of intentions and potentially harmful behaviors.

**Figure 7:**
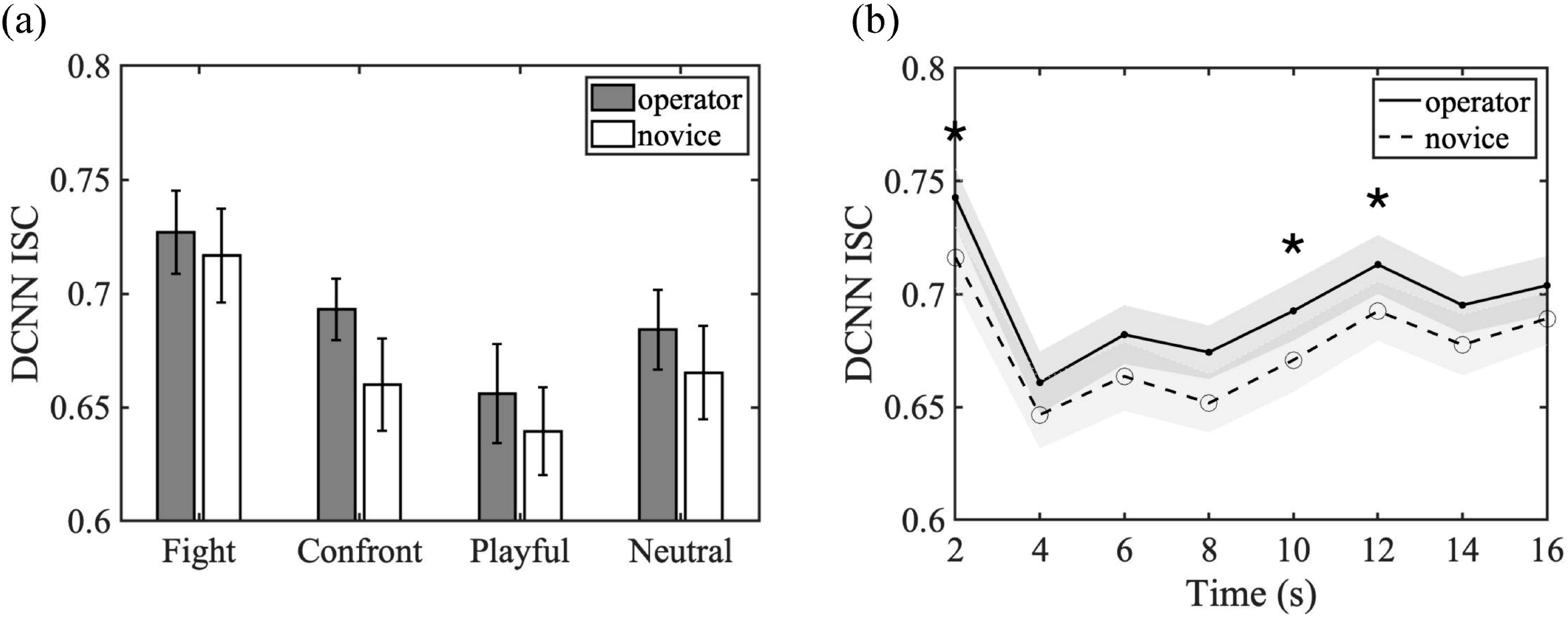
DCNN ISC results. (a) DCNN ISC of each action category as calculated by averaging pairwise correlations of DCNN FC7 features concatenated over time within the CCTV operator group or within the novice group. Error bars indicate SEs across videos in each action category. (b) Group differences in DCNN ISC over time. Shaded areas indicate SEs. Asterisks indicate significant group difference tested against a Bonferroni corrected alpha level.

## 4. Discussion

The current study adopted a saliency model and a DCNN model to examine the impact of low- and high-level visual information attended in gaze patterns of experienced CCTV operators and novices when viewing the same surveillance footage with and without harmful intentions.

For the low-level visual cues extracted by the saliency model, we did not find group-level differences in saliency indices, but classifiers based on patterns of saliency features of gaze-centered regions distinguished CCTV operators from the novices. In particular, optical-flow motion information contributed the most to the classification of the two participant groups. We also found a group difference in the ISC of saliency, showing greater consistency of using saliency information among CCTV operators. The saliency findings suggest that CCTV operators did not simply attend to regions with greater visual saliency as the overall saliency indices did not show group differences between CCTV operators and novices. However, operators attended to salient regions with a greater inter-subject correlation over time in comparison to novices, and also employed shared strategies to focus on certain patterns of visual salient cues (e.g., certain motion patterns) that likely facilitated intention inference and prediction. For the object-level features extracted by the DCNN model, we were also able to distinguish the gaze patterns of the CCTV operators from the novices using DCNN features, which contain high-level object-relevant information that reflects semantic representations of entities in visual scenes. Interestingly, the decoding performance of all action categories except neutral actions was significantly greater than chance level. This may suggest that DCNN features best distinguish CCTV operators from novices for actions with specific intentions, including fighting, confrontation, and playful events, but less well for actions with unclear intentions in the neutral condition. Additionally, CCTV operators showed higher inter-subject correlation in using similar DCNN features than novices, suggesting more similar information-seeking behavior by consistent eye movement patterns among operators when predicting potentially harmful interaction outcomes.

Here, we reliably decoded groups based on both patterns of low-level saliency features and patterns of object-relevant semantic features extracted by DCNN. These results may suggest that extensive experience of monitoring surveillance footages induces strategies in attending to different patterns of saliency and semantic features toward goal-directed actions. For example, Howard et al. (2010) found that individuals with more experience watching football matches made eye movements to goal-relevant areas of the scene earlier than non-experts. In a meta-analysis by Gegenfurtner et al. (2011), effects of expertise were robustly associated with an increased frequency of fixations on goal-relevant information and reduced latencies for first fixations on these areas. From the decoding of saliency features, we found that the most frequently used feature that distinguishes CCTV operators and novices was motion cues, which may result from efficient processing of human actions in experts. The enhanced attention to goal-relevant information (and consequently, reduced attention to irrelevant information) may underly the effect of expertise in a variety of visual tasks (Haider and Frensch, 1996). The impact of expertise on attentive features is not limited to monitoring surveillance footages, but was also revealed in art expertise (Koide, Kubo, Nishida, Shibata, & Ikeda, 2015), where the authors found that artists may extract visual information from paintings based on features such as textures and composition of colors, driven by artists’ deep aesthetic appreciation of paintings.

The higher inter-subject correlation among operators in attending to saliency features and DCNN features is consistent with previous findings about expertise in processing surveillance footages. For example, Howard et al. (2013) found that when monitoring a single scene to detect potentially suspicious events, trained CCTV operators showed greater consistency in fixation location by “knowing what to look for” compared to novices. Using the same dataset as the current study, Roffo et al. (2013) found that expert operators are more likely to focus on a small number of interesting regions, sampling them with high frequency. A neuroimaging study, using CCTV video stimuli that have some overlap with the current stimuli, also provided converging evidence by demonstrating that a different sample of CCTV operators show increased synchronization of neural responses in certain regions of the brain (i.e., bilateral anterior superior temporal gyrus, left middle temporal gyrus, left ventral striatum, and left inferior parietal lobule) than do novices (Petrini et al., 2014).

The current findings provide insight regarding what visual information is selectively attended by CCTV operators to detect harmful intentions. Previous magnetoencephalography (MEG) study showed that the recognition of human social interactions might involve different visual mechanisms than simple feedforward pattern recognition (Isik, Mynick, Pantazis, & Kanwisher, 2020). The authors found that different types of human social interactions can be decoded at around 500 ms after the onset of videos, which is substantially later than visual processing of objects, faces, emotions, gestures, and actions. For example, object pattern recognition can be decoded within 100 ms of the image onset (e.g., Carlson et al., 2013; Isik et al., 2014). Face perception elicits the signature N170 response at around 170 ms after face image onset (Bentin, Allison, Puce, Perez, & McCarthy, 1996), while many facial properties such as age, gender, and identity can be decoded even earlier (Dobs, Isik, Pantazis, & Kanwisher, 2018). Communicative gestures (Redcay & Carlson, 2015) and single-person actions can be decoded as early as 200 ms (Isik, Tacchetti, & Poggio, 2018). Thus, unlike visual processes of static visual patterns or single-agent movements, the inference of intentions from human social interactions may involve the recognition of high-level semantics and relational reasoning that go beyond visual pattern recognition.

A few limitations should be addressed in future studies. The fully connected layer of DCNN takes increasingly complex visual feature patterns extracted by a sequence of convolutional layers and develops invariant representations of objects that resemble the inferior temporal (IT) cortex (e.g., Yamins et al., 2014; Cichy, Khosla, Pantazis, Torralba, & Oliva, 2016). However, even though the AlexNet model was pre-trained to recognize 1000 object categories, it does not contain all the entities often encountered in surveillance footages. This limitation may restrain the formation of efficient representations of agents and objects, which are necessary to build up high-level semantic representation for intention inference. Future studies with DCNNs that are more specialized in video understanding and scene analysis may further advance the probe of high-level semantic information contributing the expert’s recognition of intentions in social interactions. For example, the two-stream CNN (Simonyan & Zisserman, 2014) inspired by the two-stream processing of biological motion perception in the brain provided a qualitative account of some behavioral results observed in human biological motion perception (Peng, Lee, Shu, & Lu, 2020) and may be used in future investigations.

Together, the current study combines eye movement data with computational analysis to reveal the impact of intensive training with surveillance footage on the visual processing of human interactions from a unique perspective. The results from the two computational analyses indicate that CCTV experience facilitates the detection and recognition of intention from natural videos via actively processing low-level visual saliency and object-level semantic information. Novices may be momentarily distracted by unimportant visual cues that do not necessarily inform the upcoming social outcomes. In contrast, CCTV operators may consistently and strategically direct the selective attention toward visual regions revealing goal-relevant semantics, such as a person walking toward a group of people who may end up joining the fight. Indeed, part of CCTV training includes developing awareness for a whole scene to acquire evidence about all relevant people and objects (Walker, Tyerman & Porter, 2021). The current results not only shed light on how extensive experience shapes up visual processing of complex stimuli in biological systems, but also illustrate the promise of using computational models to analyze visual information attended by different groups of participants. Furthermore, our findings imply that computer vision algorithms that incorporate both visual pattern recognition in images and semantic encoding of the inter-person relationship at the abstract level may advance the ability of AI in inferring social intentions and making predictions on harmful outcomes.

## Acknowledgments

This work was funded by the Human Dimension and Medical Research Domain of the Dstl Programme Office for the UK Ministry of Defence and NSF BSC-1655300. We thank all the CCTV staff and other participants for their contribution to this research. We thank Steven Thurman for conducting preliminary analyses for this project.

## Author Contributions

Developing the study concept: YP, FP, HL. Funding acquisition: FP, HL. Computational modeling: YP, JB. Human data collection: CN. Data analysis: YP, GT, JB. Writing – Original Draft Preparation: YP. All authors provided critical revisions and approved the final version of the paper for submission.

## Data Statement

Due to the sensitive nature of the CCTV footage used in this study, raw stimuli data will remain confidential and cannot be shared.

